# Reclassification of a likely pathogenic Dutch founder variant in KCNH2; implications of reduced penetrance

**DOI:** 10.1101/2022.08.05.502917

**Authors:** J.S. Copier, M. Bootsma, C. Ng, A.A.M. Wilde, R.A. Bertels, H. Bikker, I. Christiaans, S.N. van der Crabben, J.A. Hol, T.T. Koopmann, J. Knijnenburg, A.A.J. Lommerse, J.J. van der Smagt, C.R. Bezzina, J.I. Vandenberg, A.O. Verkerk, D.Q.C.M. Barge-Schaapveld, E.M. Lodder

## Abstract

**Background:** Variants in *KCNH2*, encoding the hERG channel which is responsible for the rapid component of the cardiac delayed rectifier K^+^ current (I_Kr_), are causal to Long QT Syndrome type 2 (LQTS2). We identified eight index patients with a new variant of unknown significance (VUS), *KCNH2*:c.2717C>T:p.(Ser906Leu). We aimed to elucidate the biophysiological effect of this variant, to enable reclassification and consequent clinical decision-making.

**Methods:** A genotype-phenotype overview of the patients and relatives was created. The biophysiological effects were assessed by manual whole-cell patch-clamp using HEK293a cells expressing: (I) wild type (WT) *KCNH2*, (II) *KCNH2*-p.S906L alone (homozygous, Hm) or (III) *KCNH2*-p.S906L in combination with WT (1:1) (heterozygous, Hz). A calibrated automated patch-clamp assay using Flp-In HEK293 was used to follow up on the functional data.

**Results:** Incomplete penetrance of LQTS2 in *KCNH2*:p.(Ser906Leu) carriers was observed. In addition, some patients were heterozygous for other VUSs in *CACNA1C, PKP2, RYR2*, or *AKAP9*. The phenotype of carriers of *KCNH2*:p.(Ser906Leu) ranged from asymptomatic to life-threatening arrhythmic events. Manual patch-clamp showed a reduced current density by 69.8%, and 60.4% in KCNH2-p.S906L-Hm and *KCNH2*-p.S906L-Hz, respectively. The time constant of activation was significantly increased with 80.1% in *KCNH2*-p.S906L-Hm compared to *KCNH2*-WT. Assessment of KCNH2-p.S906L-Hz, by calibrated automatic patch-clamp showed a reduction in current density by 35.6%.

**Conclusion:** The reduced current density in the *KCNH2*-p.S906L-Hz indicates a moderate loss of function. Combined with the reduced penetrance and variable phenotype, we conclude that *KCNH2*:p.(Ser906Leu) is a low penetrant likely pathogenic variant for LQTS2.

## INTRODUCTION

Long QT syndrome (LQTS) is a well-known cardiac arrhythmic disorder affecting 1:2000 of the population[1]. The syndrome is characterized by a prolongation of the QT segment corrected for heart rate (QTc) measured by 12-lead electrocardiography (ECG), and ECG morphology abnormalities. The prolonged QT resembles a delayed repolarization of cardiac action potentials, which can result in Torsade de Pointes (TdP). This can lead to syncope and sudden cardiac death (SCD)[2]. The syndrome can either be induced by certain drugs, ionic imbalance or be hereditary. In the latter case, a genetic cause can often be found. However, not all carriers of a genetic variant are symptomatic, meaning that there is an incomplete penetrance. To date, variants in eight genes are recognized as causal[3]. However, the evidence for causality of the pathogenic variants differs[4]. The majority of the pathogenic variants, 75%, are in *KCNQ1* (MIM:607542), *KCNH2* (MIM:152427), or *SCN5A* (MIM:600163), leading to LQTS type 1, 2, or 3, respectively[5].

The *KCNH2* gene encodes for the human ether a-go-go (hERG) channel, also known as KCNH2 or Kv11.1. The hERG channel is responsible for the rapid component of the delayed rectifier K^+^ current (I_Kr_). Genetic variants in *KCNH2* can cause a loss of function in hERG, resulting in a reduced I_Kr_ current[6]. According to the standard classifying system, the variants can be (likely) benign, (likely) pathogenic, or a variant of unknown significance (VUS)[7]. The classification of the variant impacts the clinical treatment of the patients and the genetic counselling offered to the patients and their relatives.

To prevent overdiagnosis, substantiated suspicion of LQTS, based on the Schwartz score, is needed before patients are referred for detailed genetic testing[8–11]. The Schwartz score, based on ECG parameters, medical history and family history, scores a patient as having a low, intermediate, or high probability of LQTS. Unfortunately, after genetic testing a substantial amount of VUSs are identified. VUSs with a potentially reduced penetrance present a clinical challenge as collecting definitive data concerning the effects of such variants depends on both large sample sizes and functional data. The Finnish founder variants, which show an incomplete penetrance, are a well described example of this[12]. Due to the complexity of the assessment, little attention has been devoted to these kinds of variants.

In this paper, we assess the clinical presentation of the carriers and the effect of the *KCNH2*:c.2717C>T:p.(Ser906Leu) variant on hERG functionality. This variant has been briefly mentioned by others in studies genetically assessing arrhythmia patients[13,14]. However, a detailed clinical description and segregation data is lacking, and no functional assessment of the variant has been performed. As a consequence, to date, the variant is classified as VUS, hampering the counseling of the patients and their relatives. The current study describes eight unrelated families from the Netherlands in which the *KCNH2*:p.(Ser906Leu) variant was found. We provide a detailed picture of the variable clinical presentation of the patients and assess the biophysiological properties of the mutated hERG channel, with the ultimate aim of reclassifying the *KCNH2*:p.(Ser906Leu) variant.

## MATERIALS AND METHODS

### Clinical assessment

Written informed consent was collected from all participants under research protocols (VUmc_2020_4231 and W20_226 # 20.260) approved by the local medical ethical committee. Clinical data was collected from index patients who were assessed by their cardiologist upon suspicions of LQTS. This was done independently in four Dutch academic medical centers. ECGs were recorded from all patients, in several also Holters and exercise tests were studied. ECG parameters were determined manually, and QT was corrected by the Bazetts method. In addition, LQT-related symptoms and family history was collected. Based on these findings genetic testing and counseling were offered. Following diagnoses of LQTS, clinical assessment of first-degree relatives was actively pursued. Relatives were offered genetic testing of the *KCNH2* variant in presence of a reasonable suspicion of LQTS or to assess segregation.

### Genotype analysis and segregation

The *KCNH2*-p.S906L variant has been identified in the index patients and family members by Next Generation Sequencing or Sanger sequencing, respectively, in a period ranging from 2009 to 2022. All gene panels used contained the major LQTS-related genes; *KCNQ1, KCNH2*, and *SCN5A*. Other genes often sequenced were *CACNA1C* (MIM:114205), *CALM1* (MIM:114180), *CALM2* (MIM:114182), *CALM3* (MIM:114183), *KCNE1* (MIM:176261), *KCNE2* (MIM:603796), *KCNJ2* (MIM:600681), *KCNQ1* and *TRDN* (MIM:603283). The index patients of families D, F, and G had genetic analysis by additional panels, containing a larger group of arrhythmia- and cardiomyopathy-related genes ranging from 37 to 50. A complete list of sequenced genes per patient is available in Table S1. No other potential LQTS variants were identified in the patients other than those described in the patient histories below.

### Site-directed mutagenesis and cell culture

The wild-type (WT) plasmid encoding hERG/GFPires[15] was genetically altered by mutagenesis to create the S906L-hERG/GFPires plasmid, encoding *KCNH2*, NM_000238.3:c.2717C>T:p.(Ser906Leu), rs199473435. The mutagenesis was performed with the Quickchange-XLsite-directed Mutagenesis Kit (Agilent Technologies, Santa Clara, USA) and custom-designed primers. Successful insertion of the variant was confirmed by Sanqer Sequencing. Mutagenesis and sequencing primer sequences are available upon request.

Before transfection, Human embryonic kidney (HEK-293A) cells were cultured in 6-well plates at 60-80% confluence. The culture medium consisted of DMEM (21969-035, Gibco, USA), heat-inactivated fetal bovine serum (FBS)(Biowest, France), Pen-Strep (15140-122, Gibco, USA), and L-glutamine (25030-024, Gibco, USA). Cells were cultured in an incubator ensuring a constant CO2 level of 5%, at 37°C. Followed by transfection with the HERG/GFPires (WT) or S906L-hERG/GFPires (*KCNH2*-p.S906L) construct. Transfection was performed using the Lipofectamine 2000 Transfection Reagent (11668-019, Invitrogen), according to manufacturer instructions. In brief, the transfection mix contained the lipofectamine and plasmid in a 1:3 ratio and was supplemented to the transfection medium. The transfection medium was of the same composition as the culturing medium but without Penicillin-Streptomycin and heat-inactivated FBS. Transfection was done either with WT or *KCNH2*-p.S906L alone (homozygous), for heterozygous expression the cells were transfected with WT and *KCNH2*-p.S906L-plasmid in a 1:1 ratio. Cells were transfected for 4 hours at 37°C, followed by a single washing step, after which they were further cultured until measurement. Electrophysiological measurements were performed on cells exhibiting green fluorescence 36-48 hours post-transfection. Transfection and measurements for the different experimental groups were run in parallel to each other.

### Electrophysiological measurements

#### Manual patch clamp

HERG currents were recorded at 36±0.1°C with the whole-cell patch-clamp method using an Axopatch 200 B amplifier (Molecular Devices, Sunnyvale, CA, USA). Voltage control, data acquisition, and analysis were realized with custom software. Single cells were obtained by 1.0 min of trypsinization with 0.25% Trypsin EDTA (15575-020, Ultrapure, 15090-046, ThermoFisher) after which they were put in a cell chamber located on the stage of an inverted microscope (Nikon, Eclipse T*i*) before they were superfused with modified Tyrode’s solution containing (mmol/l): 140 NaCl, 5.4 KCl, 1.8 CaCl2, 1.0 MgCl2, 5.5 glucose, 5.0 HEPES, pH was set at 7.4 with NaOH. Patch pipettes (2.0-2.5 MΩ) were pulled from borosilicate glass (Harvard Apparatus, Waterbeach, UK) and filled with a solution containing (in mmol/l): 125 K-gluconate, 20 KCl, 1 MgCl2, 10 NaCl, 5 EGTA, 5 MgATP, 10 HEPES, pH was set at 7.2 with KOH. Potentials were corrected for the calculated liquid junction potential. Cell membrane capacitance (Cm) was calculated by dividing the time constant of the decay of the capacitive transient after a −5 mV voltage step from −40 mV by the series resistance. Signals were low-pass filtered at 2 kHz, and digitized at 5 and 10 kHz for (de)activation and inactivation, respectively. Series resistance was compensated for by at least 80%.

The current density was determined by dividing the current amplitude by Cm. The activation, deactivation, and inactivation kinetics of the hERG channel were assessed using the voltage-clamp protocols described previously[15] and shown in Figures 4A, 5A and 6A. Current density was calculated by dividing the current amplitude by Cm. Activation curves were fitted using the Boltzmann equation: I/Imax = A/{1.0 + exp[(V1/2 − V)/*k*]}, to determine V1/2 (membrane potential for the half-maximal activation) and the slope factor *k* (in mV). The time course of deactivation was fitted by a double-exponential equation: I/Imax = Af × exp(−t/τf) + As × exp(−t/τs), where Af and As are the fractions of the fast and slow deactivation components, and τf and τs are the time constants of the fast and slow deactivating components, respectively. The time course of activation was fitted by the mono-exponential equation: I/Imax = A × [1 − exp(−t/τ)], where A and τ are the amplitude and time constant of the activating current.

#### Automated patch clamp

A method and protocol paper detailing the design of heterozygous *KCNH2* vector, generation of Flp-In T-rex HEK293 *KCNH2* variant cell lines, cell culture routine of heterozygous KCNH2 Flp-In HEK293 for automated patch clamp electrophysiology, quality control measures, voltage protocols and data analysis was recently published[16]. In brief, DNA plasmids co-express variant and WT *KCNH2* allele were ordered from GenScript Inc (Pistcataway, NJ, USA). These plasmids were inserted into the same place in the genome using the Flp-In recombinase technology to generate Flp-In HEK293 (Thermofisher, cat. #R78007) stable cell lines. The effect of KCNH2 variants were assessed using an automated patch-clamp electrophysiology platform (SyncroPatch 384PE, Nanion Technologies, Munich, Germany) using standard voltage protocols and recording solutions described in the method and protocol paper[16]. Current density was quantified by measuring the peak amplitude of the tail current at −50 mV, after a depolarizing step to +40 mV for 1 s. Peak tail current amplitudes were normalized to Cmto obtain current density (pA/pF) and transformed to normal distribution using a square root function before normalizing to the mean of WT from the same plate. The channel deactivation was quantified as the ratio for the decay in −50 mV current amplitude at 500ms after the peak tail current.

### Statistical analysis

Data of the manual patch-clamp are expressed as mean±SEM. Normality and equal variance assumptions were tested with the Kolmogorov–Smirnov and the Levene median test, respectively. Followed by One-way ANOVA or Two-way repeated-measures ANOVA followed by a correction for multiple testing by Bonferoni’s method. p ≤ 0.05 was considered statistically significant. Data of the automated patch-clamp are presented as mean±95% CI. The thresholds for interpreting the function of *KCNH2* variant as normal or abnormal were calibrated recently against 14 benign/likely benign controls and 23 pathogenic/likely pathogenic controls[17]. Functionally normal as BS3_supporting is defined as within 2 standard deviations (SD) from the mean of benign/likely benign controls whereas functionally abnormal as PS3_supporting or PS3_moderate is defined as between 2-4 SD or >4 SD from the mean, respectively.

## RESULTS

### Clinical description

A detailed description of all families can be found in the supplemental materials. In brief, 5 male and 3 females index patients were included, with an age range of 8 till 73 years. They were clinically assessed because of non-LQTS related cardiac complaints, recurrent syncope, TdP, or out of hospital cardiac arrest (OHCA). One of the patients later died at age 21 due to SCD. After clinical assessment, genetic testing was performed using an arrhythmia or LQTS panel (Fig. S2). One index was included in an unrelated study, here the *KCNH2-*variant was identified, after which she was included in our study. Genetic counseling was offered to the index patients, as well as clinical assessment of relatives. The QTc in index patients and relatives carrying the variant ranged from normal to prolonged, accompanied by variable phenotype severity (Fig.1-2, Table S1).

**Figure 1:**
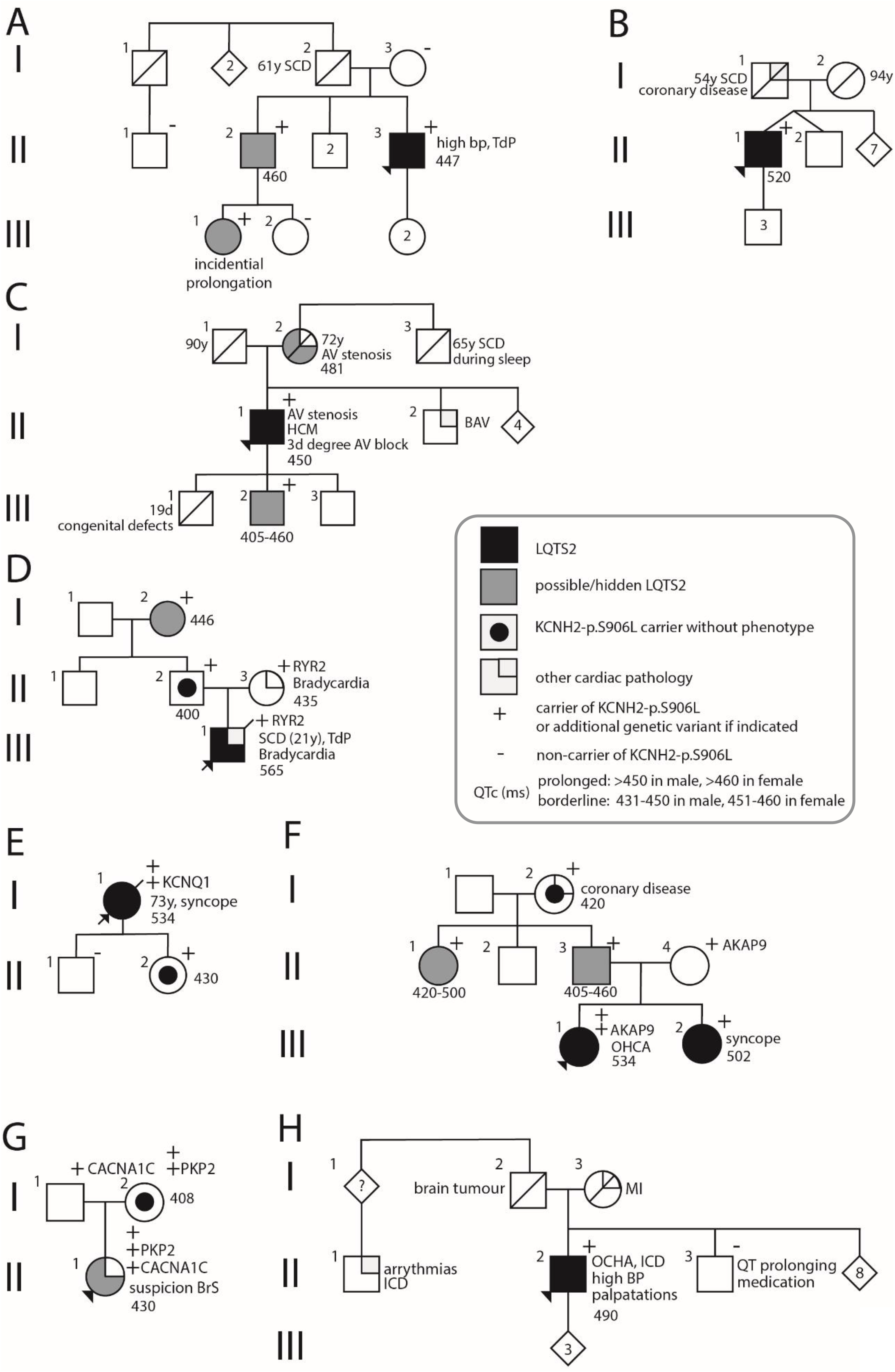
Pedigrees of index patients depicting genotype and phenotype. The *KCNQ1* variant is pathogenic (class 5), the others are of unknown significance (VUS). Phenotype is based on QTc, ECG morphology, symptoms and, if available, Holter measurements and exercise test. A clinical overview can be found in table 1. AF: atrial fibrillation, *AKAP9*: A-Kinase Anchoring Protein 9, AV: aortic valve, BAV: bicuspid aortic valve, bp: blood pressure, bpm: beats per minute, BrS: Brugada Syndrome, *CACNA1C*: Calcium Voltage-Gated Channel Subunit Alpha1 C, HF: heart failure, ICD: implantable cardioverter defibrillator, *KCNH2*: Potassium Voltage-Gated Channel Subfamily H Member 2, *KCNQ1*: Potassium Voltage-Gated Channel Subfamily Q Member 1, LVH: left ventricular hypertrophy, OHCA: out of hospital cardiac arrest, *PKP2*: plakophilin-2, QTc: QT interval corrected for heart rate, *RYR2*: ryanodine receptor 2, SCD: sudden cardiac death, TdP: Torsade de Pointes.

**Figure 2:**
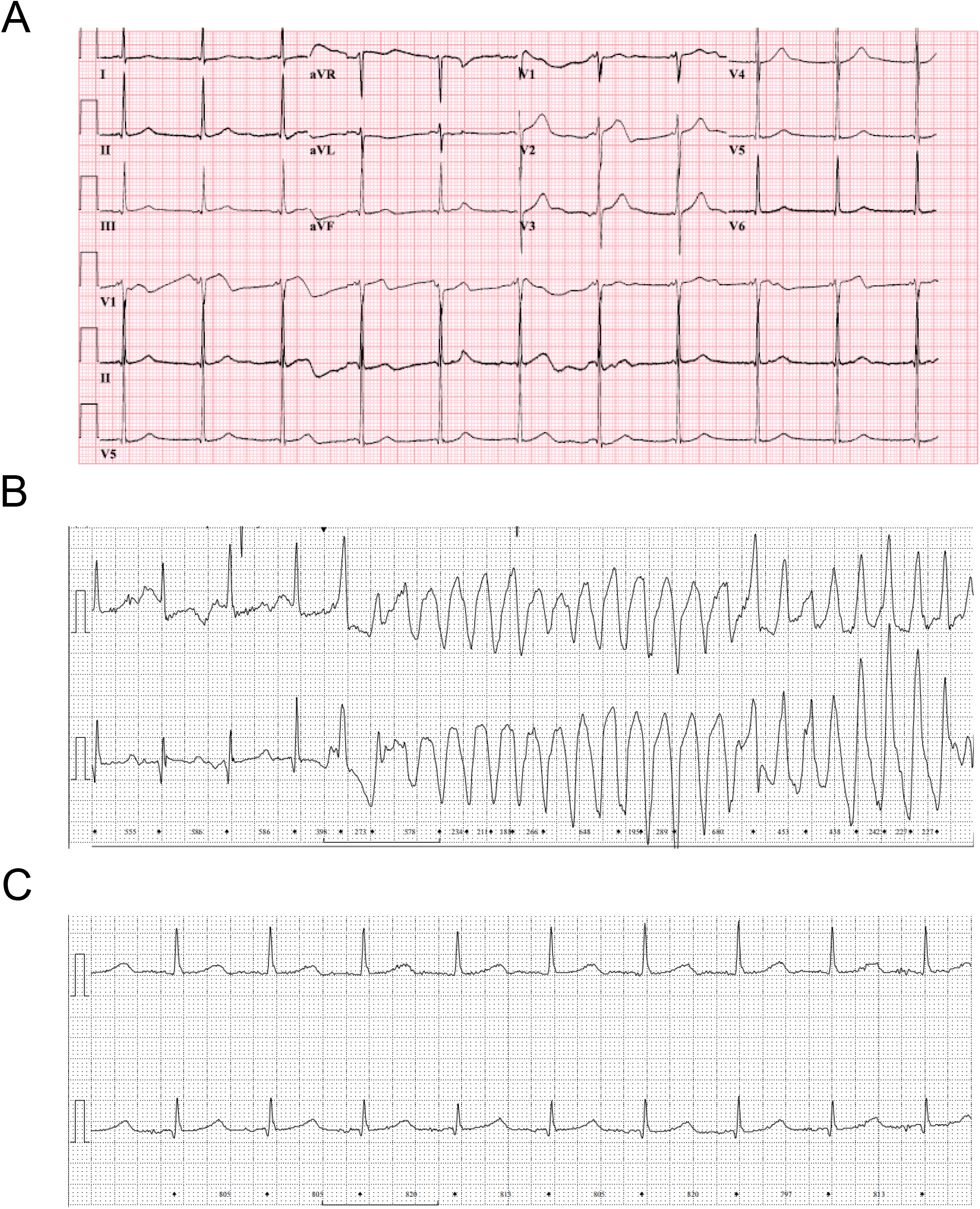
representative ECGs of index patients carrying the *KCNH2-*p.S906L variant. (A) 12-lead ECG from the index of Family C, showing a flattened ST-segment in lead I, V4 and V5, and QT prolongation (QTc 470). (B) Holter recording of Index E during a TdP episode. (C) Holter recording of index E showing QT prolongation (QTc 534).

### Genetic testing

In summary, the variant *KCNH2*:c.2717C>T:p.(Ser906Leu), was identified in all 8 index patients. Segregation analysis in family members resulted in the identification of 12 additional carriers, and 8 non-carriers. All patients diagnosed with (probable/hidden) LQTS2 who underwent genetic testing carry the variant. However, 5 currently unaffected individuals were carrying the variant. Unless otherwise described, all individuals genetically tested did not carry any other variants potentially causal for the LQTS. The identified variants were *RYR2*:c.6952A>G (MIM:180902), *AKAP9*:c.11230G>T (MIM:604001), *PKP2*:c.1114G>C (MIM:602861) and *CACNA1C*:c.6272A>G (MIM:114205), in index patients D, F, and G, respectively (Table S1).

According to the gnomAD database, the *KCNH2*:p.(Ser906Leu) variant has a MAF of 0.00004 (6/152162, gnomAD v3.1.2) in the general population, indicating that it is rare. The variant has not been detected in the GoNL cohort of Dutch references[18]. Furthermore, the amino acid concerned is highly conserved among several animal species (Fig. 3A). The variant is not predicted to be damaging based on the SIFT, Polyphen, and CADD scores, being 0.17, 0.015 and 22 respectively. The revel score does indicate it to be likely disease-causing, with a score of 0.758. The variant is located at the C-terminus of the protein, distal to the cyclic nucleotide-binding domain (CNBD)(Fig. 3B).

**Figure 3:**
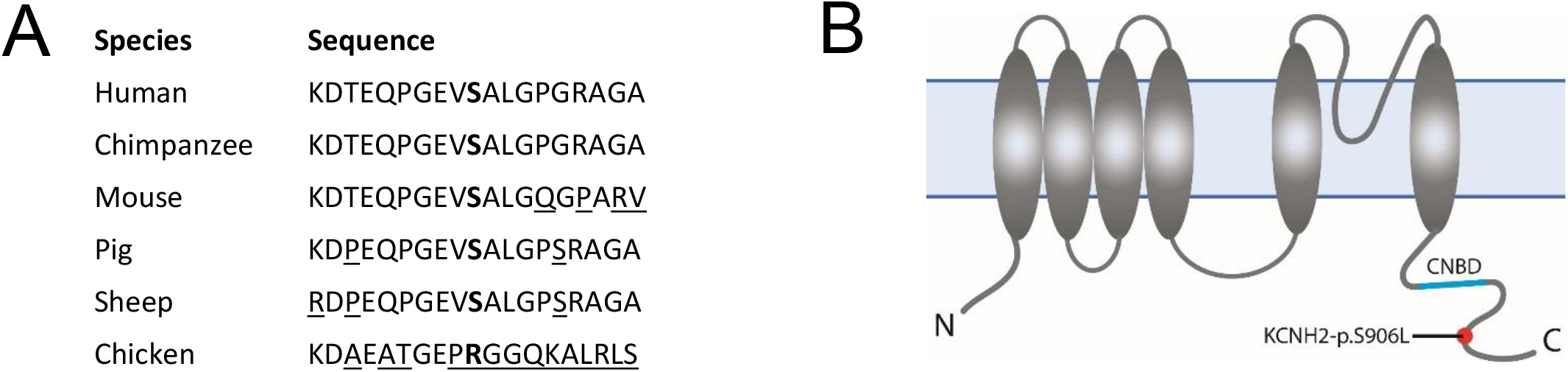
Location of amino acid substitution resulting from genetic variants in KCNH2. (A) Conservation of the Serine amino acid located at position 906 (**bold**) in *KCNH2* among animal species. (_:non-identical amino acids). (B) Localization of amino acid substitute in the KCNH2 protein. The variant is located at the distal end of the c-terminus, distal from the cyclic nucleotide binding domain (CNBD).

Haplotyping of four index patients (Fam D-G) identified a shared haplotype consisting of 291 shared SNPs encompassing a region of 929kb (928712 bp), from rs6250256 to rs150723265 (Fig. S1). This indicates that the *KCNH2:*p.Ser906Leu variant is most likely a founder mutation in the Dutch population.

### Biophysiological properties of *KCNH2*-p.S906L

Considering the diverse clinical phenotype of the carriers of *KCNH2*:p.(Ser906Leu), classification of this variant is not straightforward. We, therefore, assessed the current density and channel kinetics of the *KCNH2*-p.S906L encoded hERG by manual and automated whole-cell patch-clamp.

#### Manual whole-cell patch-clamp

##### KCNH2-p.S906L leads to a reduced current density

Steady-state and tail current densities were measured using a double pulse protocol (Fig. 4A). Steady-state currents were established by the 4 sec, P1 depolarizing pulse from a holding potential of −80 mV. These pulses result in the activation of the hERG channels, followed by the inactivation at more depolarized potentials. Typical examples and average current densities of the steady-state current are shown in Figures 4A and 4B, respectively. Current density was measured in cells expressing WT channels, or the KCNH2-p.S906L channel in a Hm or Hz manner at the steady-state (pulse 1, P1) and peak tail current (pulse 2, P2) (Fig. 4A). The steady-state current density of KCNH2-p.S906L-Hm and –Hz is significantly lower than KCNH2-WT. For example, at −30 mV, WT current density was 128.2±30.0 pA/pF, compared to 37.8±6.3 and 20.1±5.1 pA/pF in Hz and Hm, respectively (Fig. 4B, Table S2). Expressed in percentages this is a 70.5%, and 84.3% reduction, respectively. Tail current densities were analyzed from the peak currents measured during P2 in which the channels recover from the inactivation during P1, resulting in maximal availability of the channels. The density of the tail current is also significantly lower in both *KCNH2*-p.S906L models (Fig. 4C). For example, at 40 mV, KCNH2-WT tail density was 107.9±14.3 pA/pF, and 42. 8±6.2 and: 32.6±9.5 pA/pF) for Hz and Hm, respectively (Table S2). Expressed in percentages, this is a 60.4% and 69.7% reduction, respectively. Neither steady-state current nor tail current densities differs significantly between KCNH2-p.S906L-Hm and –Hz.

**Figure 4:**
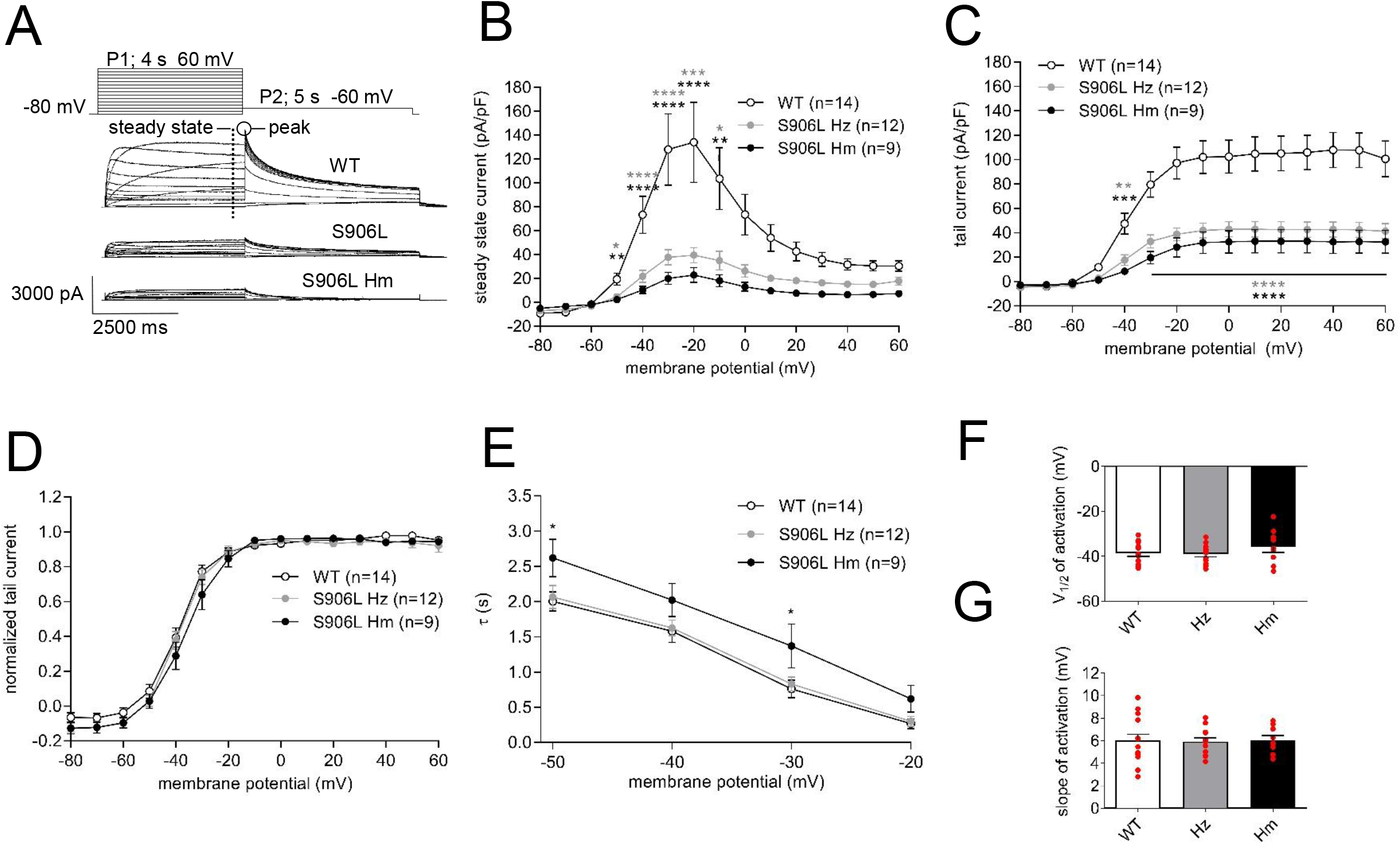
Current density and activation gating. (A) Representative traces of HEK293A cells expressing wild type (WT) KCNH2, KCNH2-p.S906L heterozygous (S906L Hz), and homozygous (S906L Hm). Inset, double-pulse protocol used. (B) Average density of steady state current. (C) Average density of peak tail current. (D) Speed of activation. (E) Voltage dependency of activation. (F and H) Half maximal activation (V _1/2_), and slope factor(*k*), determined by Boltzmann fit through activation curves. Statistics by two-way ANOVA-RM, or one-way ANOVA, and Bonferroni’s correction for multiple testing. *; p ≤ 0.05, **; p ≤ 0.01. WT; wild type, Hz; heterozygous, Hm; homozygous.

##### Minor activation gating changes as a result of KCNH2-p.S906L

From the current measured during the double pulse protocol depicted in Figure 4A, we next analyzed the voltage dependency of activation and speed of activation. For the voltage dependency of activation, we normalized the current measured during P2 of each cell to its maximal value. The obtained curves were fitted using the Boltzmann equation to obtain the V1/2 and *k*. (Fig. 4D). Neither the V1/2 (Fig. 4F) nor *k* (Fig. 4G)(Table S2) differs significantly between the groups. The speed of activation was characterized during P1 by fitting a mono-exponential equation through the currents at a voltage of −50 to −20 mV. The speed of activation is slower in KCNH2-p.S906L-Hm compared to the WT and Hz channels, while it does not differ significantly between KCNH2-p.S906L-Hz and WT (Fig. 4E). For example, at −30 mV, the τwere 0.76±0.12 (WT), 0.83±0.10 (Hz), and 1.37±0.31 (Hm) ms.

##### Minor changes in inactivation of voltage dependency and deactivation properties

In addition to activation gating, minor changes in voltage dependency of inactivation and deactivation properties in KCNH2-p.S906L-Hm compared to WT were found. Detailed description of these measurements and results can be found in the supplemental material.

#### Automated whole-cell patch-clamp

The recently calibrated automated patch clamp assay can assess the function of *KCNH2* variants to provide functional evidence for use in variant classification[17]. KCNH2-p.S906L was generated as heterozygous stable HEK293 cells and its function was assessed using this assay. The peak tail current density recorded at −50 mV for WT and KCNH2-p.S906L (Fig. 5A) shows a reduction of 35.6% in KCNH2-p.S906L compared to WT (Fig. 7B,F). KCNH2-p.S906L has a slower deactivation than benign *KCNH2* variants, serving as controls (Fig. 7C), higher steady-state channel inactivation than benign controls (Fig. 5D) and V1/2 of activation is shifted towards depolarized potential (Fig. 5E). The data points of KCNH2-p.S906L are distributed between normal and abnormal range for current density and the values for the V1/2 (Fig. 5F-G). Lastly, the channel deactivation and inactivation are functionally normal (Fig. 5H-I). Therefore, the evidence strength of KCNH2-p.S906L is indeterminate.

**Figure 5:**
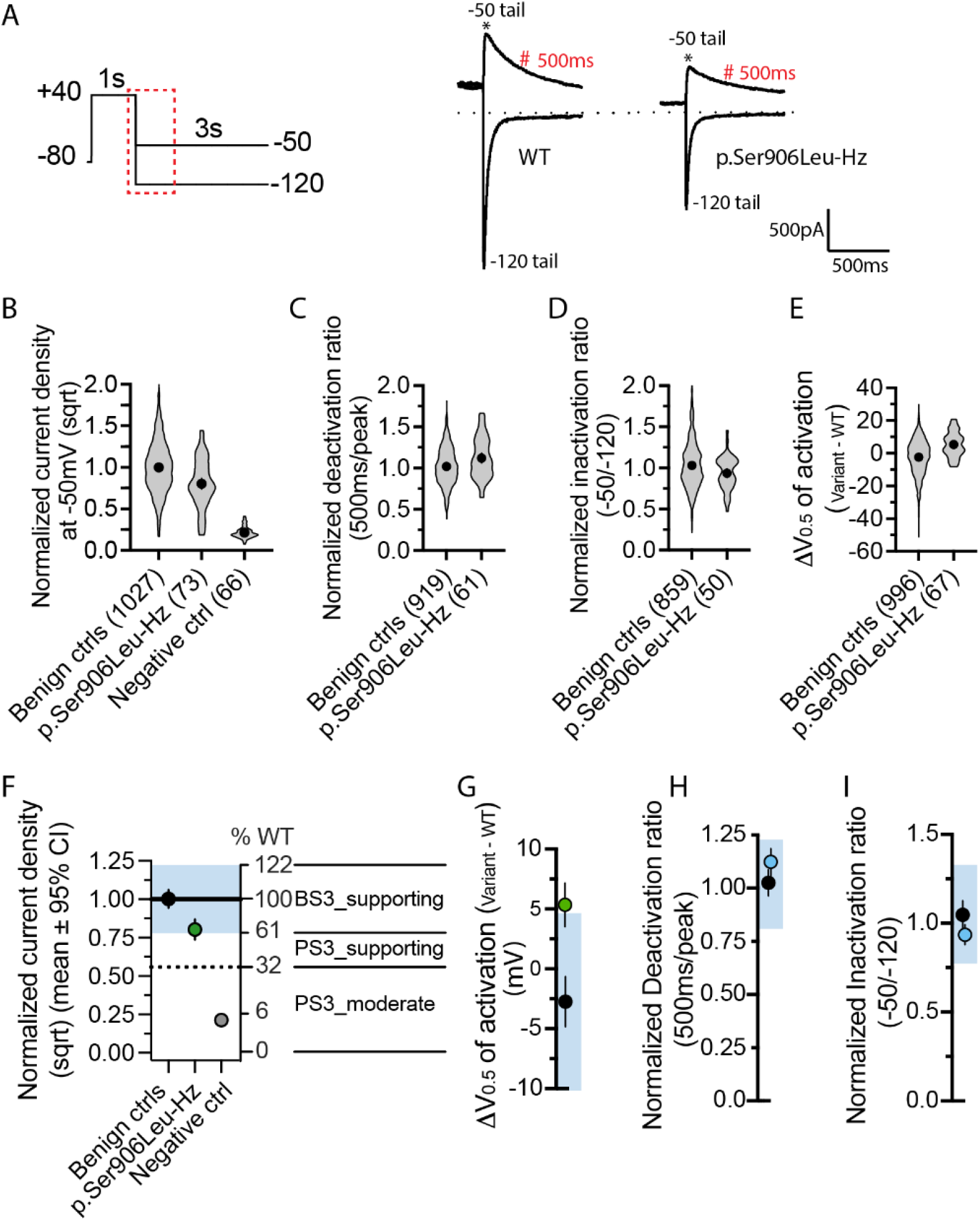
Variant interpretation using calibrated automated patch clamp assay. (A) The −50 and −120mV tail current traces corresponding to the highlighted region within the voltage protocol for WT and heterozygous p.Ser906Leu. Peak tail current and current amplitude 500ms after peak were indicated by * and #, respectively. (B) Data distribution of normalized current density (sqrt) by WT from the same plate for all the benign controls (n=1027) from Jiang et al., (2022), p.Ser906Leu (n=73 from 4 replicates) and negative control (n=66). (C) Data distribution of normalized deactivation ratio for all the benign controls (n=919) and p.Ser906Leu (n=61 from 4 replicates). (D) Data distribution of normalized inactivation ratio for all the benign controls (n=859) and p.Ser906Leu (n=50 from 4 replicates). (E) Data distribution of shift in the V_0.5_ relative from WT for all the benign controls (n=996) and p.Ser906Leu (n=67 from 4 replicates). (F) p.Ser906Leu is indeterminate for the current density as the confidence interval (CI) is spreading between functionally normal (blue region) and abnormal range. (G) p.Ser906Leu is indeterminate for shift in activation as the CI spreads between functionally normal and abnormal range. (H) p.Ser906Leu is functionally normal for channel deactivation. (I) p.Ser906Leu is functionally normal for channel inactivation. All the circles are mean± 95% CI.

## DISCUSSION

### Summary

We here describe the clinical and functional characteristics of the *KCNH2*:p.(Ser906Leu) variant. From the clinical data, the *KCNH2* variant can be regarded as a low penetrant risk factor for LQTS type 2. Assessment of the effects of the variant on the biophysiological properties of the encoded hERG channel was performed, showing a 35.6-60.4% reduction in current density measured by automated or manual patch-clamp respectively, while changes in gating properties were minor or absent. Combining the diverse clinical presentation, and the observed changes in electrophysiological properties of the channel, we conclude that this variant acts as a risk factor for LQTS2. Based on the combined classification system for variants[7] the variant can be classified as either pathogenic or likely pathogenic grounded on the weighing of the individual criteria, however with a reduced penetrance.

### Genetic heterogeneity

In the families described, it is noteworthy that multiple, severely affected patients carry additional genetic variants or have additional cardiovascular morbidities. This observation is in line with the notion that the *KCNH2*:p.(Ser906Leu) variant confers a risk towards developing LQTS2, which could be influenced by additional genetic variants as described in various studies[19–21]. Assessment of the functional effect of the additional VUSs found was beyond the scope of this study. Of note, variants *in CACNA1C* and *AKAP9* have been suggested to be associated with LQTS, however, this causality is being disputed[4,22–24]. Nonetheless, they may modify the extent of the QT prolongation[21,25,26].

In our families we observe an incomplete penetrance, there have been other studies describing this regarding *KCNH2* variants. For instance, two Finnish founder variants, R176W and L552S, with 23-35% and 29-38% of the carriers being symptomatic, respectively[12,27,28]. The QTc in the carriers are 459±40 (R176W), and 463±45 (L552S)[12]. Risk assessment showed an increased risk for cardiac events in carriers compared to non-carriers. However, this risk is not as high as in carriers of a non-founder variant[29]. Moreover, it seems that Finnish compound heterozygous carriers are more severely affected than carriers of a single variant[27]. The observed QTc values in carriers of the Finnish founder variants are on average borderline prolonged, and there might be an additive effect of second genetic variants, this is similar to our observations.

### Interpretation patch-clamp data

Our results indicated a 35.6% and 60.4% loss in current density in a heterozygous expression model, when measured by automated or manual patch-clamp respectively. This is an intermediate reduction when compared to pathogenic variants, and could therefore explain the mild phenotype in most carriers.

This is in line with the previously described *KCNH2*:p.G816V variant which was identified in mildly affected patients, and resulted in an approximate 50% reduction of current density[30]. Reports on functional studies of pathogenic variants describe a 70-90 % reduction in current density[31–33], or substantial changes in gating properties[34,35], when measured by manual patch-clamp. Application of automated patch-clamp to assess pathogenic variants, indicated that the majority have a reduction in current density of at least 75% when expressed heterozygously[36]. Recently, it has been shown that a reduction of ±39-68% is supportive of being (possibly) pathogenic[17]. Despite there being a difference between the loss of function found by manual- and automated patch-clamp in our study, both support the variant resulting in a moderate loss of function. The observed disparity in values measured by the different techniques could be due to methodological variations.

Previous studies have described *KCNH2*-variants with a disconcordance between the functional data and the clinical phenotype of the carriers[37,38]. This is possibly influenced by secondary factors like drugs, ion levels, clinical comorbidities, or additional genetic variants. The latter is substantiated by research finding an additive effect of a *KCNH2* polymorphism, combined with a causal variant in *KCNH2* on current density[38]. Furthermore, the earlier discussed Finnish founder variants do not decrease the current density but cause an increased deactivation rate[27,28]. Despite the moderate changes in channel functionality, these carriers proved to have a prolonged QTc and an increased risk of cardiac events[29]. This emphasizes that even variants with a moderate effect on channel functionality like ours can increase the risk of cardiac events.

### Risk assessment & treatment

In the clinic, cascade screening of family members is performed when a (likely) pathogenic (class 4 or 5) variant is identified in the index patient[39,40]. Based solely on the current clinical data, the *KCNH2*-p.S906L variant would not classify as (likely) pathogenic according to the ASHG criteria[7] even though multiple carriers have a clear phenotype and are at high risk for a cardiac arrhythmic event[41]. However, our electrophysiological data shows that there is a moderate reduced current density, this would add a PS3 argument to the classification resulting in a (likely) pathogenic classification. However, the current guidelines do not incorporate the option of moderate results in a functional test, hampering this classification. Based on the Finnish data and other studies, even carriers with a normal QTc are possibly at a higher risk than non-carriers[27,28,42]. Therefore, we would recommend counseling by a clinical geneticist, and the clinical and genetic assessment of the first-degree relatives of carriers of the *KCNH2*:p.(Ser906Leu) variant. Those identified to carry the variant should be clinically followed-up every 5 years. Currently, there are no official recommendations regarding the treatment and counseling of carriers of variants with a reduced penetrance such as *KCNH2*:p.(Ser906Leu). More awareness and research are needed to record these modifying genetic variants to reach a consensus regarding the treatment and counselling of these carriers. We propose the active treatment of those with a phenotype. Lifestyle accommodation, including avoidance of QT prolong drugs, are pertinent to both carriers with a phenotype and those with a concealed phenotype.

### Limitations

In our experimental design, we use a heterologous expression model by introducing human *KCNH2* into HEK293 cells, which do not express *KCNH2* themselves. In this way, we aimed to study the functionality of the hERG channel. To mimic heterozygous expression of the variant we co-transfected with the WT and KCNH2*-*p.S906L plasmid, however both have the same fluorescent reporter. Thereby there is no visual conformation of successful transfection with both plasmids. Nevertheless, we observed no values suggesting imbalanced transfection. Transfection of HEK293 cells enables the assessment of the functionality of hERG. However, this system does not mimic the effect this variant could have on more complicated systems such as cardiomyocytes or on whole heart level. In these systems, interactions between ion currents are present contributing to action potential generation and propagation, and cellular contraction.

In addition, the data used for this study are collected in retrospect. Thereby we were not able to create an entirely uniform dataset. Furthermore, not all individuals discussed underwent the same clinical assessment leading to not all having been subjected to an exercise test or Holter measurement.

Lastly, we were unable to assess if this variant is enriched compared to the general population, as the total number of LQTS patients screened for this variant is unknown. However, the low frequency of this variant in gnomAD is very suggestive[43].

## Conclusion

We conclude that *KCNH2*:p.(Ser906Leu), in combination with additional factors, is a low penetrant likely pathogenic variant for LQTS2. Carriers should be informed about the risk of QT prolonging drugs and hypokalemia, which, combined with the variant, could further increase the risk of a cardiac event[2,44,45]. Additionally, carriers should undergo regular cardiac assessment. In final, for those with a severe phenotype or early disease-onset, extending the genetic testing to the general arrhythmia panel should be considered for the detection of possible second (pathogenic) variants.

## ACKNOWLEDGEMENTS

The authors wish to thank all the patients and their relatives for participating in this study.

## CONTRIBUTORS

All authors contributed to the acquisition, analysis, or interpretation of the data. JSC was responsible for drafting the manuscript. All authors revised and approved the final manuscript.

## SUPPLEMENTAL MATERIAL

