## Supplemental Materials for "Reclassification of a likely pathogenic Dutch founder variant in KCNH2; implications of reduced penetrance"

**SUPPLEMENTAL RESULTS**

**Clinical description**

Family A

The male index (II-5), presented at age 59 years with high blood pressure and a suspected TdP which was cardioverted. ECG shows a QTc of 447 at baseline, with a near-normal QT morphology. Genetic testing revealed the *KCNH2*:p.(Ser906Leu) variant. Two family members were clinically assessed, the index's brother (II-2) is asymptomatic, and has a QTc of 460 with a normal ECG morphology. His daughter (III-1) had incidental QT prolongation detected by a 24-hours Holter measurement. Both are carriers of the variant. Family history revealed that the father of the index died aged 61 due to sudden cardiac death (SCD) (I-2). Because the asymptomatic mother of the index (II-3) did not carry the variant, he might have been an obligate carrier.

Family B

The 67-year-old male index patient (II-1) has a QTc of 520 and panel testing identified the *KCNH2*-variant. There has been no genetic testing of any of the index's relatives. None of the eight siblings have to date any known symptoms or cardiac problems, and no ECGs were available. The index patient had three sons, one died at birth due to *solutio placentae*. The other two sons have had no recorded symptoms or clinical assessment. Both parents have died, the father (I-1) aged 54 due to SCD possibly caused by coronary disease. The mother (I-2) died aged 94 without having any known cardiac problems.

Family C

The index (II-1) is part of a family with a broad history of cardiac-related pathologies. At age 66 he had a cardiac evaluation because of a bicuspid aortic valve in his brother (II-2). He was diagnosed with mild aortic valve (AV) stenosis, and LQTS with a QTc of 450 (Fig. 2A). Panel testing lead to the identification of the *KCNH2*:p.(Ser906Leu) variant. Three years later a pacemaker was implanted for a 3$^{rd}$-degree atrioventricular block (AVB). Aged 74, he collapsed due to polymorphic ventricular tachycardia (PVTs), hypertrophic cardiomyopathy (HCM) was detected, no cardiomyopathy-related variants were identified in follow-up genetic testing. One of the index's sons (III-2) has a QTc of 460 and a normal exercise test, he is carrier of the variant. There is no genetic information available regarding other family members.

Family D

At age 21 the male index (III-1) patient died due to SCD while resting. Prior to the event, a loop recording was implanted because of previously experienced syncope and bradycardia. The patient died before the results were analyzed. These revealed a QTc of 565 and subsequent TdP leading to SCD. DNA-diagnostics of arrhythmia genes identified, the paternally inherited *KCNH2*:p.(Ser906Leu) variant as well as an maternally inherited VUS in *RYR2:*c.6952A>G, which has a mean allele frequency (MAF) of 0.00001 (gnomAD, v3.1.2.). The index's father (II-2), has experienced a single episode of syncope. However, cardiac investigation showed no LQTS-related abnormalities. The mother of the father (I-2), also carrier of the *KCNH2*-variant, has a borderline LQTS2 phenotype. The index's mother has no LQTS-related abnormalities, she does have bradycardia with a heart rate (HR) of 55 beats per minute (bpm). Her father (I-V), a carrier of the *RYR2*-variant, experienced two syncopal events aged approximately 40 years old and was known to have a very low heart rate prior to developing atrial fibrillation (AF).

Family E

The index patient concerns a 73-year-old female (I-3) with recurrent syncopal events. ECG analysis showed a QTc of 534 and abnormalities in QT morphology fitting LQTS. Holter monitoring confirms a clear QT prolongation and TdP was evidenced, confirming the diagnosis (Fig 2B-C). Genetic testing identified a known pathogenic variant in *KCNQ1*:c.1760C>T, and the *KCNH2*:p.(Ser906Leu) variant. The index's daughter (II-2), carrier of only the *KCNH2*-variant, has a QTc of 430 and has experienced one syncope.

Family F

Aged 18 years old the female index patient (III-1) suffered an out-of-hospital cardiac arrest (OHCA) evoked by ventricular fibrillation. Clinical assessment showed a QTc of 524, and abnormalities in ECG morphology at baseline and during exercise test. All findings fit the LQTS2 diagnosis, and genetic testing identified the *KCNH2*:p.(Ser906Leu) variant, inherited from her father. In addition, a maternally inherited VUS in *AKAP9*:c.11230G>T, with a MAF of 0.00005 (gnomAD, v3.1.2.) was identified. Her sister (III-2) experienced recurrent near-syncopal events. At age 19, she had a QTc of 502 and was diagnosed with LQTS. The index's father (II-3) is asymptomatic but reports some cardiac discomfort at the beginning of exertion, he has a QTc of 405-460. The father's sister (II-1) is asymptomatic and has a QTc of 420. The father's mother (I-2) had a normal QTc and suffers from coronary disease. All discussed family members carry the *KCNH2*-variant. As of yet, only the mother of the index (II-4) has been identified as carrier of the *AKAP9*-variant, she has a normal phenotype.

Family G

The index patient has a phenotype of an arrhythmogenic disease, which does not completely fit with LQTS. She underwent genetic testing as part of an unrelated research project in one of the medical centers contributing to this study. The genetic test identified *KCNH2*:p.(Ser906Leu), resulting in her inclusion in this study. She was clinically assessed at 8 years old and had a QTc of 430 at baseline ECGs, which is within normal limits[18]. After experiencing premature ventricular contractions (PVC), she received flecainide treatment. ECGs taken during this treatment were indicative of Brugada syndrome (BrS) Type 1. Both her parents (I-2, I-3) are asymptomatic and have a normal QTc, baseline ECG and exercise test. The lack of phenotype in her parents precludes segregation analysis. Additional genetic screening with a broad panel (suppl. Table 1) identified the following variants: *CACNA1C*:c.6272A>G, and *PKP2*:c.1114G>C (MAF of 0.007362 (gnomAD, v3.1.2.), and 0.00069 (ALFA) respectively). Reports regarding the pathogenicity of the these variants are conflicting, as both are reported as VUS or likely benign (ClinVar). Genetic testing of the index's parents identified the mother to be carrier of the *KCNH2*- and *PKP2*-variants, and the father to be carrier of the *CACNA1C*-variant.

Family H

The index patient (II-2) experienced an OHCA at age 57, during mild hypokalemia, which was followed by the implantation of a cardioverter-defibrillator (ICD). Furthermore, he presented with hypertension and has experienced palpitations. ECG analysis revealed a QTc of 490 which remained prolonged after cessation of sotalol. Genetic panel testing identified *KCNH2*:p.(Ser906Leu). His brother (II-3) has a QTc of 480 while using QT-prolonging medication, and did not report any symptoms indicative of LQTS. He tested negative for the *KCNH2*-variant. Anamnestic, a first-degree cousin of the index patient (II-1) has a clinical history of arrhythmias, with ICD implantation, his clinical record was not available to us. Other family members had normal cardiac evaluations and were thus not genetically tested.

**Biophysiological properties of *KCNH2*-p.S906L**

Minor changes in inactivation of voltage dependency as a result of KCNH2-p.S906L

The voltage dependency of inactivation was measured using a three-pulse protocol (Fig. S2-A). The first pulse (P1) is a depolarizing pulse from -80 to 60 mV and fully activates and subsequently inactivates the current. During the short second repolarizing pulse (P2), the channels can recover from inactivation and then the voltage dependency of inactivation can be characterized by the peak of the current during the third pulse (P3). Figure S2-B shows the average inactivation curves of the three groups. The curves are virtually mostly overlapping suggesting no or minor changes in voltage dependency of inactivation. Indeed, $V_{1/2}$ (Fig. S2-C) did not differ significantly and a change in *k* was only observed in the Hm compared to the WT (WT: -21.7±1.2 mV, Hm: -30.2±5.4 mV)(Table S3).

Minor changes in deactivation properties as a result of KCNH2-p.S906L

Finally, we characterized the current-voltage (I-V) relationships of the fully activated current as well as the deactivation properties using a double-pulse protocol (Fig. S3-A). The first pulse (P1) served to activate and inactivate the current; the peak tail currents during the second pulse (P2) were used to determine the fully activated current over a large voltage range. The I-V relationships, normalized to the largest peak current, showed no significant changes between WT and *KCNH2*-p.S906L-Hz. However, a significant difference was observed between WT and Hm with larger currents at voltages ≤ -80 mV (at -80 mV; WT: 0.17±0.06, Hz: -0.04±0.04, Hm: -0.09±0.08)(Fig. S3-B, Table S3). The reversal potentials were -79.29±0.23 mV and -77.94±0.53 mV, in Hz and Hm respectively, which is significantly different to the WT (-83.30±0.28 mV). Time course of deactivation was analyzed using bi-exponential fits and did not reveal any changes in cells expressing the mutant heterozygously or homozygously, in neither the fast- nor slow component of deactivation (Fig. S3-C, Table S3). In addition, no changes were found in the contribution of the slow component to the deactivation ($A_s/(A_s+A_f)$)(Table S3).

### SUPPLEMENTAL TABLES

| Patient | | Sex | Age | QTc | ECG morphology | Type 2 | Holter | Exercise test | Symptoms | Phenotype | Genotype |
|---|---|---|---|---|---|---|---|---|---|---|---|
| A | II-5* | M | 71 | 447 | Minor deviations | +/- | NA | NA | Hypertension, TdP | + | *KCNH2:*c.2717C>T: p.(Ser906Leu)[1] |
| | II-2 | M | 56 | 460 | nl | +/- | nl | nl | none | +/- | *KCNH2:*c.2717C>T: p.(Ser906Leu)[1] |
| | III-1 | F | 21 | NA | nl | NA | incidential QT prolongation | nl | none | +/- | *KCNH2:*c.2717C>T: p.(Ser906Leu)[1] |
| | I-2 | M | 61 | NA | NA | NA | NA | NA | SCD, 61y | NA | NA |
| B | II-1* | M | 67 | 520 | NA | NA | NA | NA | NA | + | *KCNH2:*c.2717C>T: p.(Ser906Leu)[1] |
| C | II-1* | M | 66 | 450 | low volt ST segment | + | post pause accentuation | pronounced QTc prolongation | AV stenosis, HCM, 3d degree AV block | + | *KCNH2:*c.2717C>T: p.(Ser906Leu)[1] |
| | I-1 | M | 73 | 430 | NA | NA | NA | NA | Angina pectoris, MI (73y), died 90y | NA | NA |
| | I-2 | F | 72 | 481 | NA | NA | NA | NA | Coronary disease, AV stenosis, died 72y shortly after AV surgery | +/- | NA |
| | I-3 | M | 65 | NA | NA | NA | NA | NA | SCD during sleep, 65y | NA | NA |
| | II-2 | M | NA | NA | NA | NA | NA | NA | BAV | NA | NA |
| | III-2 | M | 45 | 405 | NA | NA | clear QT prolongation (QTc 460) | nl | none | +/- | *KCNH2:*c.2717C>T: p.(Ser906Leu)[1] |
| D | III-1* | M | 24 | 565 | NA | + | pause dependent ectopy, TdP | NA | SCD at 21y, TdP, bradycardia | + | *KCNH2:*c.2717C>T: p.(Ser906Leu)[1] *RYR2:*c.6952A>G: p.(Asn2318Asp)[2] |
| | I-2 | F | 79 | 446 | flat ST, clear U | +/- | no QT prolongation | nl | none | +/- | *KCNH2:*c.2717C>T: p.(Ser906Leu)[1] |
| | II-2 | M | 52 | 400 | nl, clear U | - | too long QTc during exercise. No clear post pause accentuation | occasionally longer | none | - | *KCNH2:*c.2717C>T: p.(Ser906Leu)[1] |
| | II-3 | F | 54 | 435 | nl, clear U | - | no QT prolongation | nl | Bradycardia, 55 bpm | - | *RYR2:*c.6952A>G: p.(Asn2318Asp)[2] |
| E | I-1* | F | 73 | 534 | Terminal negative. Post pause accentuatio n. Flat ST after PM implant | + | Evident LQT, TdP | clearly abnormal | Syncope, died 73y | + | *KCNH2:*c.2717C>T: p.(Ser906Leu)[1] *KCNQ1:c.1760C>T: p.(Thr587Met)*[3] |
| | II-2 | F | 44 | 430 | Flat, low QRS volt | - | nl | nl | none | - | *KCNH2:*c.2717C>T: p.(Ser906Leu)[1] |
| F | III-1* | F | 23 | 524 | flat ST, clear U | + | NA | bas prol, bigemini VES | OHCA | + | *KCNH2:*c.2717C>T: p.(Ser906Leu)[1] *AKAP9:*c.11230G>T: p.(Gly3744Trp)[4] |

| | | | | | | | | | | | |
|---|---|---|---|---|---|---|---|---|---|---|---|
| | III-2 | F | 19 | 502 | nl, clear U | + | post pause accentuation | no prolonged QTc | Syncope | + | *KCNH2:*c.2717C>T: p.(Ser906Leu)[1] |
| | II-3 | M | 57 | 405-460 | Flat ST, clear U | +/- | post pause accentuation | no prolonged QTc | none | +/- | *KCNH2:*c.2717C>T: p.(Ser906Leu)[1] |
| | II-4 | F | 54 | 410 | nl | - | nl | nl | none | - | *AKAP9:*c.11230G>T*:* p.(Gly3744Trp)[4] |
| | II-1 | F | 58 | 420 | nl | - | flat ST-T segment , QTc: 500 | no prolonged QTc | none | +/- | *KCNH2:*c.2717C>T: p.(Ser906Leu)[1] |
| | I-2 | F | 85 | 437 | Flat ST | - | NA | nl | Coronary disease | - | *KCNH2:*c.2717C>T: p.(Ser906Leu)[1] |
| G | II-1* | F | 11 | 430 | nl, T2 small | - | 2nd ectopy | no prolonged QTc | Suspicion brugada | +/- | *KCNH2:*c.2717C>T: p.(Ser906Leu)[1] *PKP2*:c.1114G>C :p.(Ala371Pro)[5] *CACNA1C*:c.6272A> G: p.(Asn2091Ser)[6] |
| | I-2 | F | 41 | 408 | nl | - | QTc: 450 | nl | none | - | *KCNH2:*c.2717C>T: p.(Ser906Leu)[1] *PKP2*:c.1114G>C: p.(Ala371Pro)[5] |
| | I-3 | M | 48 | 390 | nl | - | unknown | nl | none | - | *CACNA1C*:c.6272A> G:p.(Asn2091Ser)[6] |
| H | II-2* | M | 56 | 490 | nl | +/- | NA | nl | Hypertension, palpatations | + | *KCNH2:*c.2717C>T:p .(Ser906Leu)[1] |
| | II-3 | M | NA | 480 | nl | - | nl | nl | Using QT-prolonging medication | - | negative |
| | II-1 | M | NA | NA | NA | NA | NA | NA | Arrhythmias, ICD | NA | NA |

**Table S1: patient characteristics.** The QTc depicted are baseline values, a QTc of >450 ms in males, and >460 ms in females is considered prolonged. There are carriers with a normal QTc, but with other ECG characteristics fitting LQTS2; flat ST, clear U, abnormal exercise test. Depicted age is at the moment of last clinical assessment. Bradycardia was defined as a heart rate < 60 bpm. AF: atrial fibrillation, AV: aortic valve, BAV: bicuspid aortic valve, bp: blood pressure, bpm: beats per minute, BrS: Brugada Syndrome, F: female, HF: heart failure, ICD:cardioverter defibrillator, HCM: Hypertrophic cardiomyopathy, M: male, NA: not available, nl: normal, OHCA: out of hospital cardiac arrest, QTc: QT interval corrected for heart rate, SCD: sudden cardiac death, TdP: Torsade de pointes. [1] NM_000238.3:c.2717C>T, [2] NM_001035.2:c.6952A>G, [3] NM_000218.2:c.1760C>T, [4] NM_005751.4:c.11230G>A, [5] NM_001005242.2:c.1114G>A, [6] NM_000719.6:c.6272A>G

| Index of Family | Genes |
|---|---|
| A | *KCNQ1, KCNH2, SCN5A* |
| B | KCNQ1, KCNH2, SCN5A, KCNE1, KCNE2, MLPA |
| C | CACNA1C, CALM1, CALM2, CALM3, KCNE1, KCNE2, KCNH2, KCNJ2, KCNQ1, SCN5A, TRDN |
| D | *ABCC9, AKAP9, ANK2, CACNA1C, CALM1, CALM2, CALM3, CASQ2, CAV3, DPP6, GJA5, HCN4, JPH2, KCNA5, KCND3, KCNE1, KCNE2, KCNH2, KCNJ2, KCNQ1, LAMP2, LMNA, MYL4, NKX2-5, NPPA, PKP2, PLN, PRKAG2, RANGRF, RYR2, SCN5A, SNTA1, TECRL, TNNI3K, TNNT2, TRDN, TRPM4* |
| E | *KCNQ1, KCNH2, SCN5A* |
| F | ABCC9, AKAP9, ANK2, CACNA1C, CALM1, CALM2, CALM3, CASQ2, DPP6 (c.-340), GJA5, HCN4, HEY2, HOOK3, JPH2, KCNA5, KCND3, KCNE1, KCNE2, KCNH2, KCNJ2, KCNQ1, LAMP2, LMNA, MYL4, NKX2-5, NPPA, PKP2, PLN, PRKAG2, RANGRF, RRAD, RYR2, SCN5A, SLC4A3, SNTA1, TCAP, TECRL, TNNI3K, TNNT2, TRDN, TRPM4. |
| G | ABCC9, AKAP9, ANK2, ASPH, CACNA1C, CACNA1D, CACNA2D1, CACNB2, CALM1, CALM2, CALM3, CASQ2, CAV3, DPP6 (c.-340), GJA5, GNB2, GPD1L, HCN4, JPH2, KCNA5, KCND3, KCNE1, KCNE5, KCNE2, KCNE3, KCNH2, KCNJ2, KCNJ5, KCNJ8, KCNQ1, LAMP2, LMNA, MYL4, NKX2-5, NPPA, PKP2, PLN, PPA2, PRKAG2, RANGRF, RYR2, SCN1B, SCN2B, SCN3B, SCN4B, SCN5A, SCN10A, SLMAP, SNTA1, TECRL, TNNI3K, TNNT2,TRDN,TRPM4 |
| H | *KCNQ1, KCNH2, SCN5A, KCNE1, KCNE2* |

**Table S2: overview of genetic panels per index.** All panels at least contain the three major LQTS genes; *KCNQ1, KCNH2, SCN5A* (underlined). The number of genes sequences ranged from three to fifty.

| Channel kinetic | | *KCNH2* WT | *KCNH2*-p.S906L Hz | *KCNH2*-p.S906L Hm |
|---|---|---|---|---|
| Current density | Steady state (pA/pF) (at -30 mV) | 128.2 ±30.0 (n=14) | 37.8±6.3 *** (n=12) | 20.1±5.1 *** (n=9) |
| | Peak Tail (pA/pF) (at 40 mV) | 107.9±14.3 (n=14) | 42.7±6.2 ** (n=12) | 32.6±9.5 *** (n=9) |
| Activation | Time component ($\tau$) (s) (at -30 mV) | 0.76±0.12 (n=14) | 0.83±0.10 (n=12) | 1.37±0.31 * (n=9) |
| | $V_{1/2}$ (mV) | -38.7±1.3 (n=14) | -39.0±1.2 (n=12) | -35.7±2.6 (n=9) |
| | *K* (mV) | 6.0±0.6 (n=14) | 5.9±0.3 (n=12) | 6.1±0.4 (n=9) |
| Deactivation | I-V relation (norm) (at -80 mv) | 0.17±0.06 (n=10) | -0.04±0.04 (n=6) | -0.09±0.08 (n=6) |
| | Erev (mV) | -83.3±0.3 (n=10) | -79.3±0.2 **** (n=6) | -77.9±0.5 **** (n=6) |
| | $A_s/(A_s+A_f)$ (at -80 mv) | 0.40±0.03 (n=10) | 0.31±0.04 (n=6) | 0.42±0.03 (n=6) |
| | Time component - Fast ($\tau$) (s) (at -80 mV) | 0.12±0.02 (n=10) | 0.12±0.02 (n=6) | 0.35±0.13 (n=6) |
| | Time component - slow ($\tau$) (s) (at -80 mV) | 0.89±0.12 (n=10) | 0.77±0.14 (n=6) | 1.07±0.40 (n=6) |
| Inactivation | $V_{1/2}$ (mV) | -37.6±6.0 (n=16) | -33.9±3.3 (n=9) | -42.6±7.2 (n=6) |
| | *K* (mV) | -21.6±1.2 (n=16) | -18.7±1.5 (n=9) | -30.2±5.4 * (n=6) |

**Table S3: overview electrophysiological values measured by whole-cell patch clamp on transfected HEK293A cells.** Data is presented as mean±SEM. Statistics by two-way ANOVA-RM, or one-way ANOVA, and Bonferroni's correction for multiple testing. * ; p ≤ 0.05, ** ; p ≤ 0.01. *** ; p ≤ 0.005, ****; p ≤ 0.001 all versus WT. WT; wild type *KCNH2*, Hz; *KCNH2*-p.S906L heterozygous, Hm; *KCNH2*-p.S906L homozygous

### SUPPLEMENTAL FIGURES

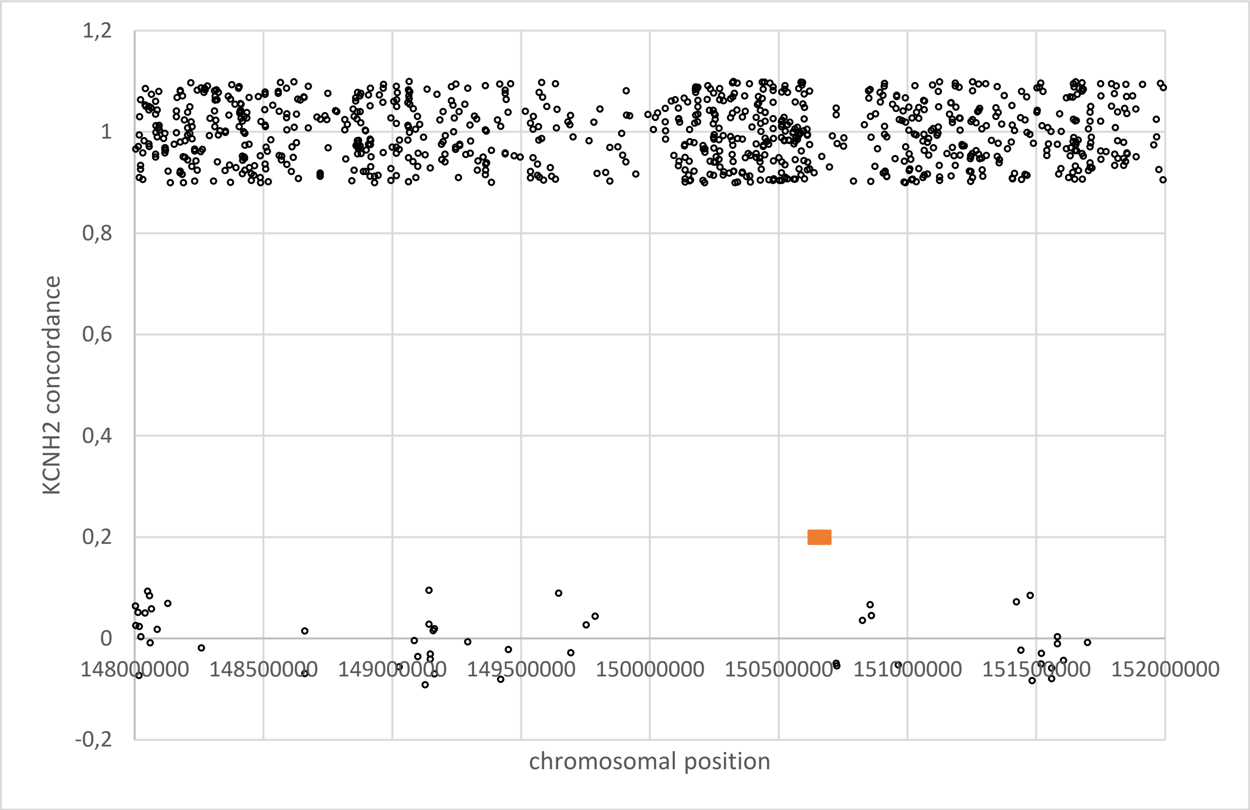


**Figure S1: haplotyping of index patients D, E, F and G by single nucleotide polymorphism (SNP) array.** Concordance between the four index patients in a section of *KCNH2* covering 40 kb. A value of ±1.0 indicates concordance, a value of ±0.0 indicates disconcordance. From location 149787501 to 150723467 (929 kb) there is a section with complete concordance, containing 291 shared SNPs.

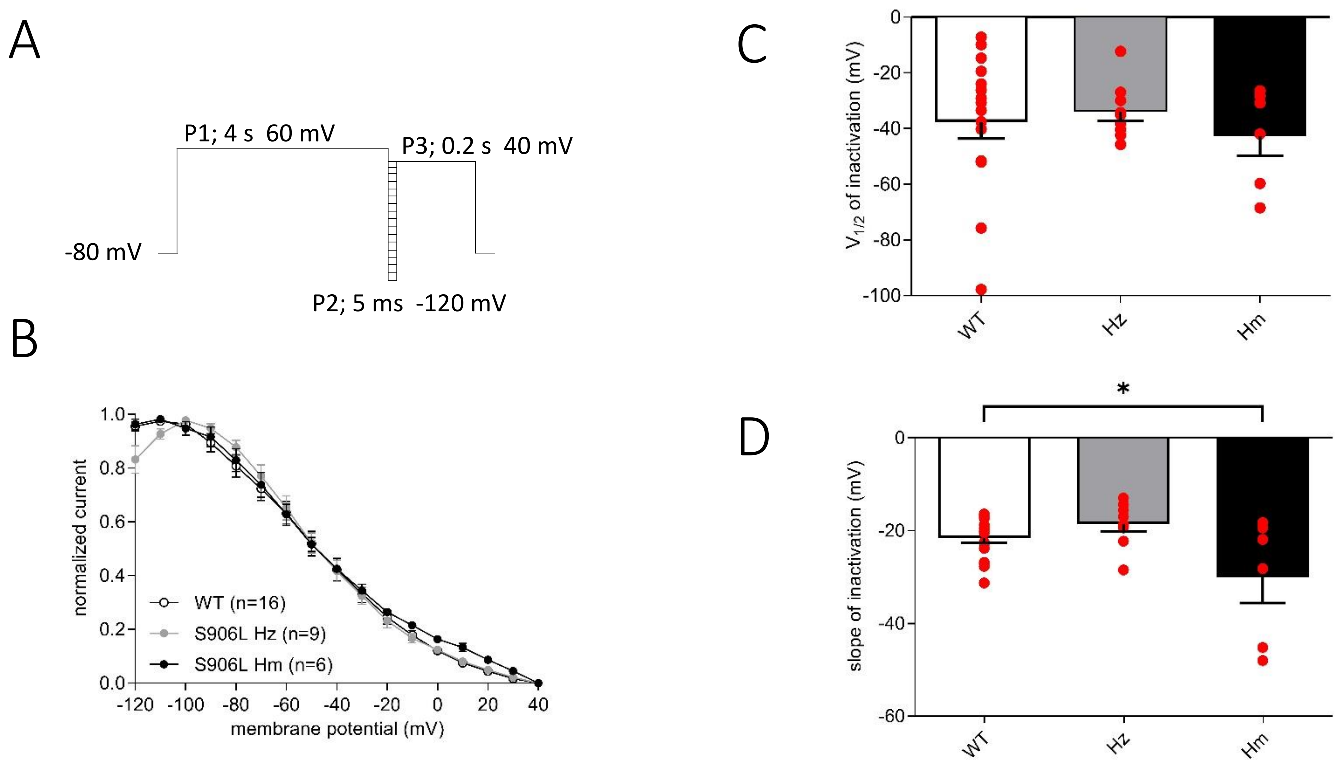


**Figure S2: Voltage dependency of inactivation.** (A) Three-pulse voltage clamp protocol applied in whole-cell patch-clamp of transfected HEK293A cells (B) Voltage dependency of inactivation. (C and D) $V_{½}$ (C), and *k* (D), determined by Boltzmann fit through inactivation curves. Statistics by two-way ANOVA-RM, or one-way ANOVA, and Bonferroni's correction for multiple testing. * ; $p \leq 0.05$. WT; *KCNH2* wild type, Hz; *KCNH2*-p.S906L heterozygous, Hm; *KCNH2*-p.S906L homozygous.

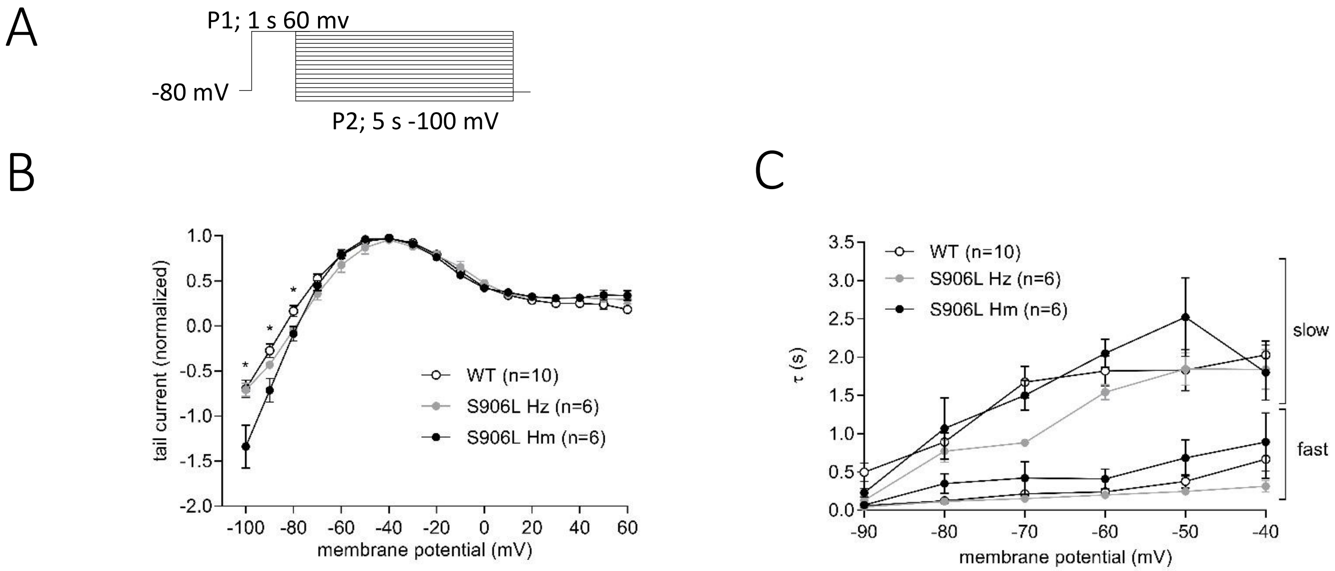


**Figure S3: Deactivation properties:** (A) Two-pulse Voltage clamp protocol applied in whole-cell patch-clamp of transfected HEK293A cells. (B) Average fully activated current-voltage (I-V) relationship normalized to maximal tail current. (C) Deactivation time course analyzed with bi-exponentional fits. Statistics by two-way ANOVA-RM, and Bonferroni's correction for multiple testing. * ; $p \leq 0.05$, WT; *KCNH2* wild type, Hz; *KCNH2*-p.S906L heterozygous, Hm; *KCNH2*-p.S906L homozygous.
